# Muc2 mucin limits *Listeria monocytogenes* dissemination and modulates its population dynamics

**DOI:** 10.1101/2020.10.21.348896

**Authors:** Ting Zhang, Jumpei Sasabe, Brandon Sit, Matthew K. Waldor

## Abstract

The mucin Muc2 is a major constituent of the mucus layer that covers the intestinal epithelium and creates a barrier between epithelial cells and luminal commensal or pathogenic microorganisms. The Gram-positive food-borne pathogen *Listeria monocytogenes* can cause enteritis and also disseminate from the intestine to give rise to systemic disease. *L. monocytogenes* can bind to intestinal Muc2, but the influence of the Muc2 mucin barrier on *L. monocytogenes* intestinal colonization and systemic dissemination has not been explored. Here, we used an orogastric *L. monocytogenes* infection model to investigate the role of Muc2 in host defense against *L. monocytogenes*. Compared to wild-type mice, we found that Muc2^-/-^ mice exhibited heightened susceptibility to orogastric challenge with *L. monocytogenes*, with higher mortality, elevated colonic pathology, and increased pathogen burdens in both the intestinal tract and distal organs. In contrast, *L. monocytogenes* burdens were equivalent in wild-type and Muc2^-/-^ animals when the pathogen was administered intraperitoneally, suggesting that systemic immune defects do not explain the heightened pathogen dissemination observed with oral infection route. Using a barcoded *L. monocytogenes* library to measure intra-host pathogen population dynamics, we found that Muc2^-/-^ animals had larger pathogen founding population sizes in the intestine and distal sites than observed in wild-type animals. Comparisons of barcode frequencies revealed that, in the absence of Muc2, the colon becomes the major source for seeding the internal organs. Together, our findings reveal that Muc2 limits *L. monocytogenes* dissemination from the intestinal tract and modulates its population dynamics during infection.

## Introduction

Muc2 is a highly abundant O-glycosylated mucin glycoprotein that is primarily found on the mucosal surface of the intestinal tract and forms a gel-like structure that is the principal component of the mucus layer found at the interface between the intestinal epithelium and lumen (1, 2). Muc2 is synthesized by goblet cells, where it is oligomerized during intracellular trafficking and stored in secretory granules prior to its secretion (3). Although distributed throughout the intestinal tract, the density and structural organization of Muc2 within the mucus layer varies between sites; for example, two thick layers of Muc2 (a ‘loose’ outer layer and a ‘firm’ inner layer) are found in the colon, while only a porous mucus layer is found in the small intestine (1, 4).

A major physiological role of Muc2 is the creation of a physical barrier that segregates the gut microbiota from the intestinal epithelium (5). This barrier function is augmented by the wire mesh-like structure of Muc2 that serves as a scaffold for binding and displaying host-derived antimicrobial peptides and microbial binding proteins (e.g. human β-defensins, Relm-β, and Zg16) (6–8). The extensive O-glycosylation of Muc2 exerts both microbe- and host-directed effects that support intestinal homeostasis, by supplying nutrients (e.g. carbohydrate moieties) to promote the expansion of gut commensal species, and by delivering tolerogenic signals to lamina propria-resident dendritic cells (9, 10). The regulation of Muc2 production and secretion by goblet cells is also integrated into intestinal defense systems against enteric pathogens. For instance, Birchenough *et al*. found that a subpopulation of goblet cells in the colon release Muc2-containing granules in response to direct sensing of pathogen associated molecular patterns via the intrinsic Nlrp6 (NOD-like receptor family pyrin domain–containing 6)-dependent inflammasome (11). In addition, goblet cells undergo hyperplasia and increase their mucus granule sizes in response to signals generated by other intestinal sentinel cells (e.g. Tuft cells) during infection (12).

The impacts of Muc2 on gut homeostasis and host defense against enteric pathogens have been revealed by studies of mice harboring a targeted knockout of the *Muc2* gene (Muc2^-/-^) (13). These Muc2^-/-^ animals exhibit a reduced gap between luminal commensal bacteria and the intestinal epithelium (5, 14), epithelial hyperplasia (13, 14), increased colonic immune cell infiltration (15), altered microbiota (16), and elevated frequencies of colon cancer (13). Similar phenotypes have also been observed in a mouse strain (the “Winnie mouse”) with a missense mutation in the *Muc2* gene (17, 18). Besides intestinal tract anomalies, Muc2^-/-^ mice have systemic inflammation, higher titers of antibodies against bacterial lipopolysaccharide (LPS) and flagellin, and elevated levels of iron in circulation (19). In addition, Muc2^-/-^ mice have been found to be more susceptible to challenges with enteric pathogens, including *Citrobacter rodentium, Salmonella typhimurium*, the nematode parasite *Trichuris muris*, and the protozoal parasite *Entamoeba histolytica* (14, 20–22).

*L. monocytogenes* is a Gram-positive foodborne bacterial pathogen that can cause enteritis as well as additional disease manifestations, such as meningitis, that result from its systemic dissemination from the intestinal tract (23). Observations from a rat ligated ileal loop model revealed that *L. monocytogenes* forms aggregates on intestinal mucus and induces goblet cell degranulation (24). Several *L. monocytogenes* surface proteins have been reported to bind to Muc2 (25), potentially mediating the pathogen’s attachment to mucus. Using the mucin-expressing cell line HT29X, Coconnier *et al*. found that listeriolysin O (LLO) is the major bacterial component that triggers mucin exocytosis (26, 27). Although goblet cell degranulation is generally recognized as a host defense strategy, *L. monocytogenes* is thought to target these mucin-secreting cells to gain access to its host receptor in the intestine, E-cadherin (28).

Here, we used Muc2^-/-^ mice and an orogastric *L. monocytogenes* infection model to investigate the role of the intestinal mucus layer in host defense against *L. monocytogenes*. In comparison to wild-type (WT) mice, Muc2-deficient animals had heightened susceptibility to orogastric challenge with *L. monocytogenes*, exhibiting elevated mortality, more severe colonic pathology and increased bacterial burden in the intestine and distal organs. In contrast, pathogen burdens were similar in Muc2^+/+^ and Muc2^-/-^ animals after intraperitoneal inoculation of *L. monocytogenes*. By using barcoded *L. monocytogenes* (29), we investigated the impact of Muc2 mucin on *L. monocytogenes* population dynamics during infection. Muc2^-/-^ animals had larger pathogen founding population sizes at intestinal and distal sites and the genetic relatedness between these bacterial populations exceeded that in WT mice. Together, these findings suggest that Muc2 guards against *L. monocytogenes* dissemination from the intestine and demonstrate that this mucin modulates pathogen population dynamics during infection.

## Results

### Muc2^-/-^ mice have heightened susceptibility to orogastric challenge with *L. monocytogenes*

To assess the role of Muc2 in host defense against *L. monocytogenes*, littermate offspring of Muc2^+/-^ breeders were orogastrically challenged with *L. monocytogenes* 10403S InlA^m^, a ‘murinized’ variant of a human clinical isolate that contains a InlA allele with enhanced binding to murine E-cadherin (30). Remarkably, 92% (11 out of 12) of Muc2^-/-^ mice died during the 8-day observation period post pathogen challenge (Fig 1a), whereas only 17% (2 out of 12) of WT (Muc2^+/+^) mice succumbed (Fig 1a). In addition, Muc2^-/-^ mice began to die earlier than WT mice (Fig 1a) and lost more weight (Fig 1b). Several of the Muc2^-/-^ mice developed diarrhea ~3 days post inoculation (DPI) (Fig S1); in contrast, WT mice do not develop diarrhea in this model. Together these observations indicate that Muc2 provides protection from the morbidity and mortality of *L. monocytogenes* infection in mice.

**Figure 1.**
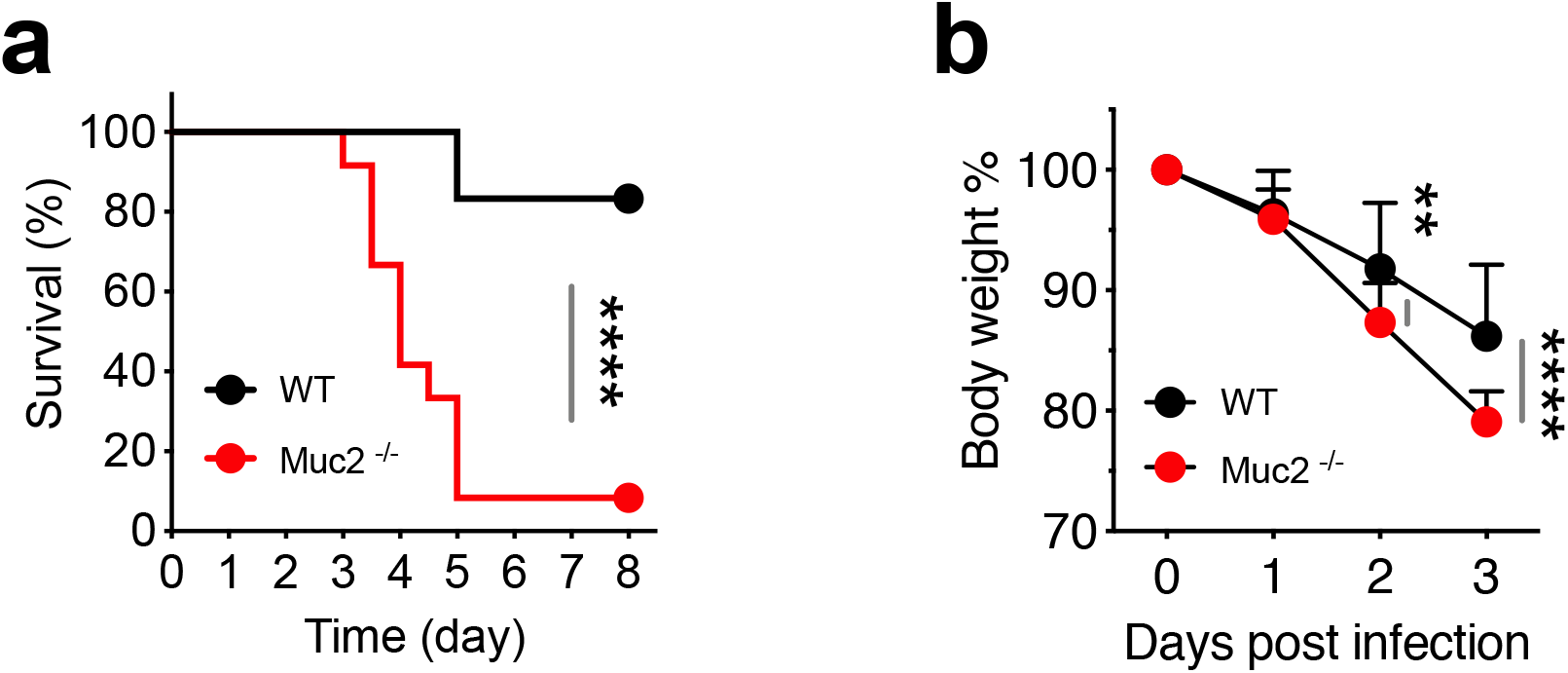
Muc2^-/-^ mice have heightened susceptibility to orogastric challenge with *L. monocytogenes*. (a) Survival curves of Muc2^-/-^ mice (n=12) and WT littermates (Muc2^+/+^, n = 12) following orogastric challenge with ~3 × 10^9^ CFU *L. monocytogenes* 10403S InlA^m^. Mice were monitored every 12 hours for 8 days. The data are from two independent experiments; the Gehan-Breslow-Wilcoxon test was used to compare the survival curves (**** p<0.0001). (b) Weight of Muc2^-/-^ (n=14) and WT (n=15) mice during the first 3 days post orogastric inoculation of *L. monocytogenes*. Data are from three independent experiments; Two-way ANOVA and Bonferroni’s multiple comparison test were used to assess significance (** p<0.01 and **** p<0.0001).

### *L. monocytogenes* infection exacerbates colonic inflammation in Muc2^-/-^ mice

The absence of Muc2 is thought to largely eliminate the physical barrier between luminal commensal bacteria and the colonic epithelium, triggering microbially-induced intestinal inflammation (15). Colons from *L. monocytogenes-infected* Muc2^-/-^ mice were visibly more swollen than colons from infected WT mice, suggestive of exacerbated intestinal inflammation in the Muc2^-/-^ group (Fig 2a, b). Furthermore, the masses of the distal colons from both uninfected and infected Muc2^-/-^ mice were greater than those from corresponding WT mice (Fig 2c); such differences were not observed in the proximal colons from either uninfected or infected animals (Fig S2). The elevated mass of the distal colon in Muc2^-/-^ mice is likely attributable to the influx of immune cells and proliferation of local colonic epithelial cells (14), but the factors that restrict these processes to the distal colon are not clear.

**Figure 2.**
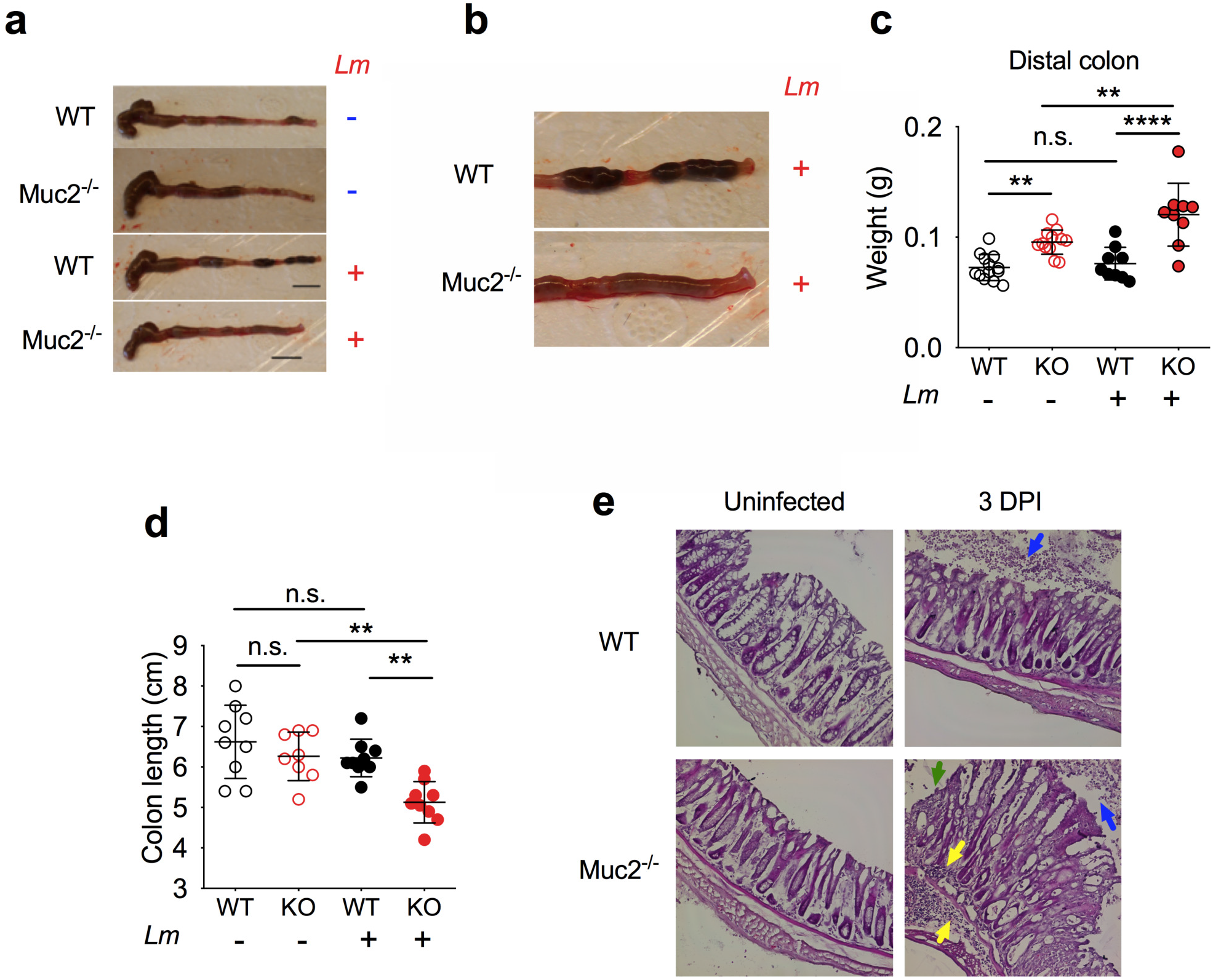
*L. monocytogenes* infection exacerbates colonic inflammation in Muc2^-/-^ mice. (a) Representative images of colons from uninfected (Lm -) and infected (Lm +) Muc2^-/-^ and WT mice at 3 days post inoculation (DPI). (b) Expanded images of distal colons shown in (a). (c-d) Weight of the distal colon (c) and length of entire colon (d) in uninfected and infected Muc2^-/-^ and WT mice at 3 DPI; 9 to 13 mice per group in (c) and 8 to 9 mice per group in (d). (e) Hematoxylin and eosin (H&E)-stained distal colonic tissue from uninfected and infected Muc2^-/-^ and WT mice at 3 DPI; the yellow arrows show immune cell infiltration and the green arrow shows sites of epithelial cell erosion; the blue arrows show granulocytes in the lumen. ANOVA and Fisher multiple comparison test were used to assess significance in c-d (n.s. not significant, ** p<0.01 and **** p<0.0001).

Inflammation of the colon is often associated with reduced colon length (31) and the colons of Muc2^-/-^ mice were significantly shorter than those from WT mice at 3 DPI (Fig 2a, d). In contrast, colon lengths were similar in uninfected Muc2^-/-^ and WT animals (Fig 2a, d), suggesting that lack of Muc2 mucin by itself is not sufficient to provoke longitudinal colon shrinkage. As *L. monocytogenes* challenge did not alter colon length in WT mice (Fig 2d), these observations suggest that infection-induced colon shortening in Muc2^-/-^ mice is due to the combined effects of the Muc2 mucin deficit and the presence of the pathogen.

Histological analysis of distal colons from WT and Muc2^-/-^ mice using H&E staining corroborated our visual observations of gross pathology. Notably, in Muc2^-/-^ mice, infection induced massive immune cell infiltration into the colonic lamina propria, heightened erosion of colonic epithelial cells, and increased granulocytes in the colon lumen (Fig 2e). In marked contrast, there was no detectable immune cell infiltration within the colonic lamina propria of WT mice (Fig 2e). Overall, with the exception of granulocytes in the lumen of the distal colon in a subset of the WT mice, there was limited pathology observed in colons from WT mice 3DPI (Fig 2e). Thus, the absence of Muc2 elevates the intestinal inflammatory response to *L. monocytogenes*.

### Muc2 mucin modulates *L. monocytogenes* intestinal colonization and dissemination

To address the impact of Muc2 on bacterial colonization, bacterial burdens in the intestines of infected animals were assessed at 3 DPI. Prior to plating, the intestinal samples were treated with gentamicin, an antibiotic that kills extracellular bacteria. In the small intestine, proximal colon, and distal colon, the Muc2^-/-^ mice carried significantly more *L. monocytogenes* than WT mice (Fig 3a-c). The most dramatic difference was observed in the distal colon, where Muc2^-/-^ mice harbored ~1000 times more *L. monocytogenes* than WT mice (Fig 3c). Thus, the absence of Muc2-containing mucin in the distal colon markedly augments the accessibility of intracellular niches to the pathogen at this site.

**Figure 3.**
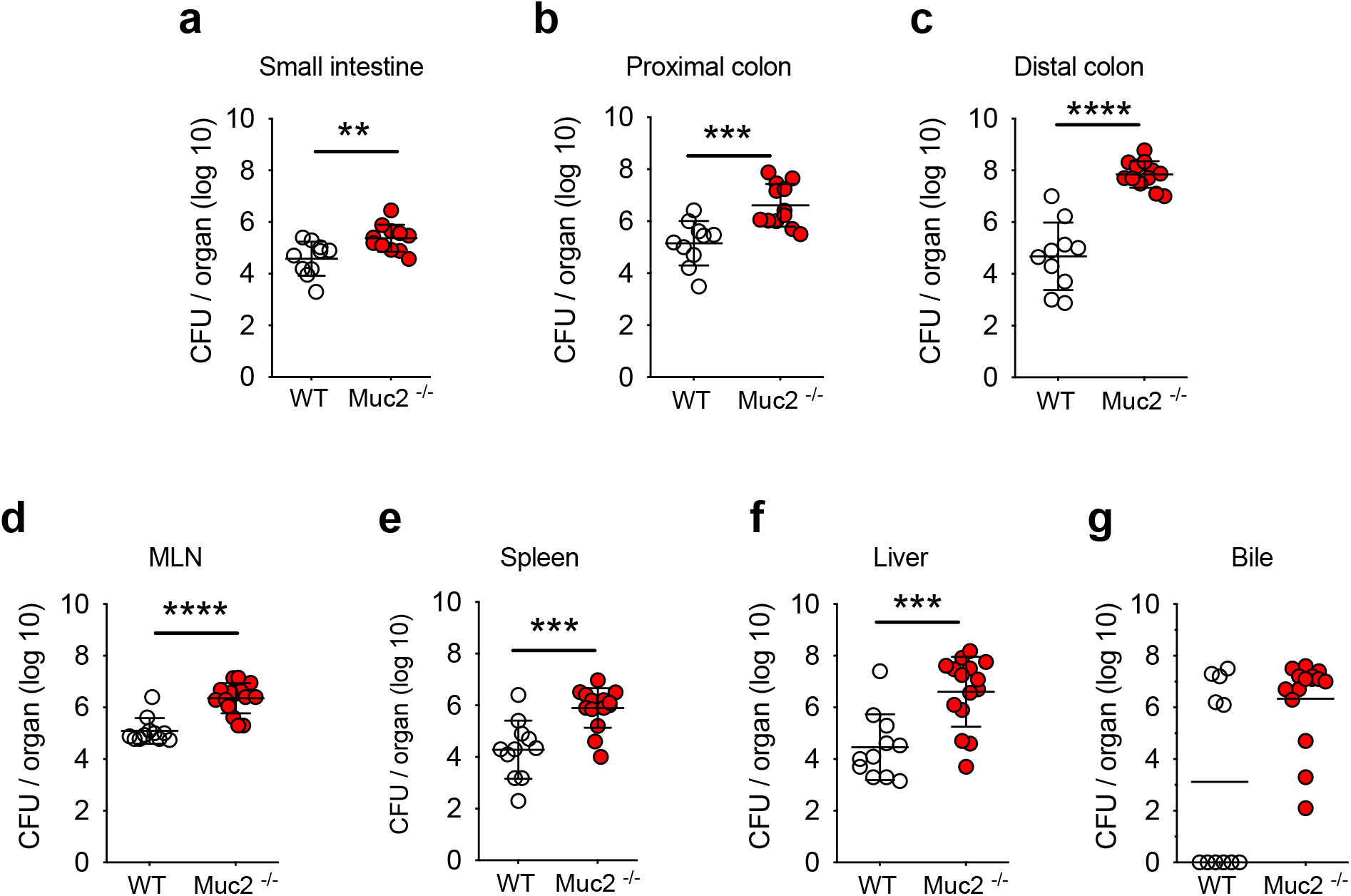
Elevated *L. monocytogenes* intestinal colonization and dissemination in Muc2^-/-^ mice. Burden of *L. monocytogenes* in the ileum (a), proximal colon (b), distal colon (c), MLN (d), spleen (e), liver (f), and gallbladder (g) at 3 DPI in orogastrically infected Muc2^-/-^ and WT mice. The intestinal samples (a-c) were treated with gentamicin to kill extracellular bacteria. Data are from three independent experiments with 10 to 15 mice per group. The Mann-Whitney test were used to assess significance (** p<0.01, *** p<0.001, **** p<0.0001).

The burden of *L. monocytogenes* in the mesenteric lymph nodes (MLN), spleen, and liver were also determined to assess the impact of Muc2 deficiency on systemic pathogen dissemination. Muc2^-/-^ mice had ~100 times higher *L. monocytogenes* burdens in MLNs, spleens, and livers than WT mice at 3 DPI (Fig 3d-f), suggesting that Muc2 contributes to the barrier that ordinarily limits *L. monocytogenes* dissemination from the gut to distal sites. Furthermore, a higher percentage of Muc2^-/-^ mice carried *L. monocytogenes* in bile recovered from the gallbladder (GB) compared to WT mice 3DPI (100 % vs 45 %) (Fig 3g). Our unpublished observations suggest that there is a positive correlation between hepatic *L. monocytogenes* burden and GB colonization, raising the possibility that the elevated frequency of GB colonization in the Muc2-deficient animals is a consequence of the higher pathogen burden in their livers. Even though the Muc2-deficient animals were more likely to have GB colonization than WT mice, the GB colonization burdens in WT and Muc2^-/-^ were similar (~10^7^ CFU, Fig 3g), suggesting that Muc2 does not alter the maximum bacterial carrying capacity of the GB.

Kumar *et al*. recently reported that Muc2^-/-^ mice have an elevated basal level of systemic inflammation (19), suggesting that the absence of a Muc2 barrier alters physiology in distal organs. To test whether Muc2^-/-^ mice have deficiencies in controlling systemic *L. monocytogenes* infection, WT and Muc2^-/-^ mice were challenged with 10^5^ CFU of *L. monocytogenes* via the intraperitoneal (i.p.) route. In contrast to our observations with oral inoculation (Fig 3), similar numbers of *L. monocytogenes* were recovered from the MLN, spleen, and liver in WT and Muc2^-/-^ mice 3 days following i.p. inoculation (Fig 4a-c), suggesting that absence of Muc2 mucin does not compromise the capacity of these organs to control the infection. In addition, Muc2^-/-^ and WT mice exhibited similar weight loss and had comparable colon lengths post i.p. challenge (Fig 4d-e), suggesting that the susceptibility of Muc2^-/-^ mice to oral pathogen challenge is not explained by a systemic immune defect.

**Figure 4.**
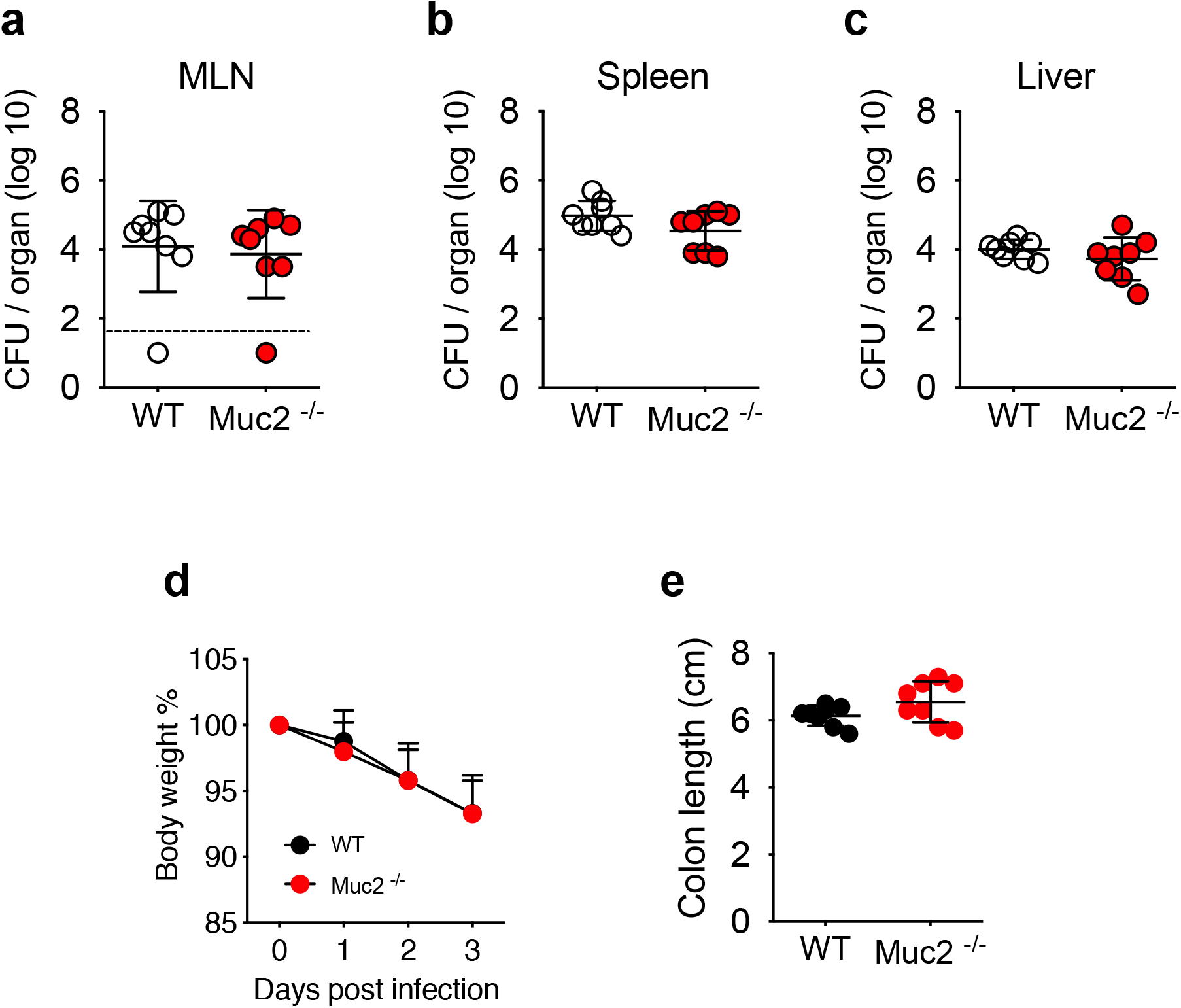
Muc2^-/-^ and WT littermate mice have similar pathogen burdens and disease manifestations post intraperitoneal challenge with *L. monocytogenes*. Muc2^-/-^ (n=8) and WT (n=8) mice were challenged with 1 × 10^5^ CFU of *L. monocytogenes* 10403S InlA^m^ via the intraperitoneal route. Pathogen burdens recovered from MLN (a), spleen (b), and liver (c) at 3 DPI. (d) Weight of Muc2^-/-^ and WT mice for three days following i.p. inoculation. (e) Colon lengths were measured at 3 DPI.

### Muc2 mucin alters *L. monocytogenes* population dynamics

Since we found that the Muc2-containing intestinal mucus barrier modulates *L. monocytogenes* intestinal colonization and dissemination, we leveraged Sequence Tag-based Analysis of Microbial Populations (STAMP) (29, 32) to investigate the impact of Muc2 on *L. monocytogenes* population dynamics during infection. In this method, DNA-barcoded, but otherwise WT, *L. monocytogenes* are used to calculate the number of bacteria from the inoculum that seed various infection sites (the founding population, (*N*b)) (29). Notably, in orally infected mice, *N*b values from intracellular (gentamicin-treated) proximal and distal colon samples were significantly higher in Muc2^-/-^ vs WT animals (Fig 5a, b). The difference was particularly pronounced in the distal colon where *N*b was ~2000 in Muc2^-/-^ vs ~50 in WT mice (Fig 5b), suggesting that Muc2 provides an especially stringent barrier for the pathogen at this site. Presumably, the absence of the Muc2 mucin facilitates pathogen access to permissive niches within colonic epithelial cells. In addition, the increased influx of immune cells and their accumulation in the colonic lamina propria of Muc2^-/-^ mice (Fig 2e) may provide additional niches for *L. monocytogenes* growth, since the pathogen is known to replicate in these cells (33).

**Figure 5.**
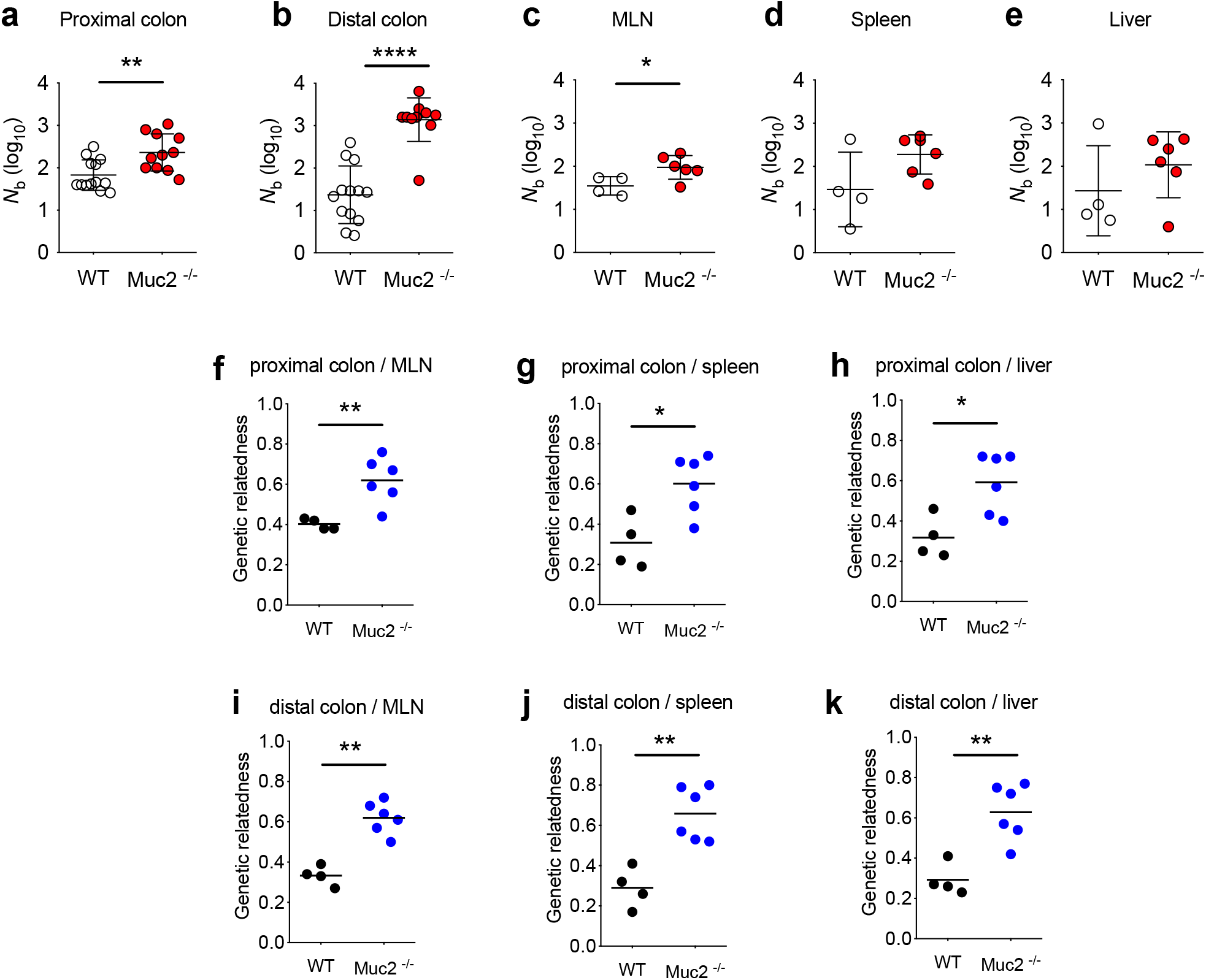
*L. monocytogenes* population dynamics differ in Muc2^-/-^ and WT mice. Founding population size (*N*b) of *L. monocytogenes* recovered from proximal colon (a) and distal colon (b) of Muc2^-/-^ (n = 11) and WT (n =13) mice at 3 DPI; data are from three independent experiments. *N*b of *L. monocytogenes* recovered from MLN (c), spleen (d), and liver (e) of Muc2^-/-^ (n = 6) and WT (n =4) mice at 3 DPI. (f-k) Genetic relatedness of *L. monocytogenes* recovered from different sites of Muc2^-/-^ (n = 6) and WT (n =4) mice 3 DPI. Mann-Whitney tests were used to assess significance in a-k (* = p<0.05, ** p<0.01, and **** p<0.0001).

*N*b sizes in internal organs also differed between Muc2^-/-^ and WT animals. *N*b values in MLNs, spleens, and livers of Muc2^-/-^ mice were higher than in WT mice, but only reached statistical significance in MLNs (Fig. 5c-e). These observations are consistent with the idea that the Muc2 mucin barrier imposes a bottleneck on *L. monocytogenes* dissemination from the intestine. In contrast, even though 100% of Muc2^-/-^ mice harbored *L. monocytogenes* in their GBs, *N*b sizes in GBs from Muc2^-/-^ were similarly extremely low (mean *N*b = 3, Fig. S3) as observed before (29), suggesting that Muc2 does not contribute to the host barrier that restricts *L. monocytogenes* access to the GB.

By calculating changes in the relative frequency of barcodes, STAMP also enables comparison of the genetic relatedness between bacterial populations recovered from different sites (32). In WT mice, bacterial populations from the proximal (Fig 5f-h) or distal (Fig 5i-k) colon were relatively distinct from those resident in the MLN, spleen, and liver (i.e. small ‘genetic relatedness’ values). These low values are similar to observations we made with Balb/c mice (29), which are more permissive to *L. monocytogenes* infection than C57BL/6 mice, and indicate that *L. monocytogenes* disseminates from the intestinal tract to distal organs using multiple independent routes (29, 34). In contrast, in Muc2^-/-^ mice, these comparisons revealed a statistically significant higher degree of relatedness between bacterial populations recovered from either the proximal (Fig 5f-h) or distal (Fig 5i-k) colon and those from the MLN, spleen, and liver than found in WT animals. These observations suggest that the increases in absolute and founding *L. monocytogenes* population sizes in the proximal and distal colon (Fig 3bc and 5ab) that are associated with the absence of Muc2 mucin also enable the colon to become the major source for seeding the internal organs. Collectively, these findings reveal that the absence of Muc2 mucin alters *L. monocytogenes* population dynamics during infection.

## Discussion

Here, we found that Muc2^-/-^ mice have heightened susceptibility to orogastric challenge with *L. monocytogenes*. Compared to WT mice, animals lacking Muc2 exhibited elevated mortality, more severe colonic pathology and increased pathogen burdens in the intestine as well as in distal organs following oral inoculation of *L. monocytogenes*. The heightened sensitivity of Muc2^-/-^ mice appears to be dependent on the route of infection, since we found that *L. monocytogenes* burdens were equivalent in WT and Muc2-deficient animals when the pathogen was administered intraperitoneally. Furthermore, our experiments with barcoded *L. monocytogenes* demonstrated that Muc2 restricts *L. monocytogenes* founding population sizes, particularly in the colon. In the absence of the Muc2 barrier, the colon becomes the dominant site from which the pathogen disseminates to distal organs. Together these observations reveal that Muc2 mucin modulates *L. monocytogenes* colonization, dissemination and population dynamics.

Several defects in Muc2^-/-^ mice likely contribute to their susceptibility to orogastric challenge with *L. monocytogenes*. First, Muc2 is the dominant mucin component of the mucus barrier that physically limits the access of commensal as well as pathogenic microorganisms to the epithelial surface of the intestine (5, 35); a thinned/absent mucus layer that lacks Muc2 likely facilitates the pathogen’s capacity to approach and ultimately invade intestinal epithelial cells. This host defense mechanism has been postulated for other enteric pathogens including *Citrobacter rodentium* and *Salmonella typhimurium*, where increased direct contact of these pathogens with the colonic epithelium of Muc2^-/-^ mice was observed (14, 21). Second, increased mucus flow during enteric infection expels pathogens from the epithelial surface and is thought to be a host defense mechanism (36). Consistent with this idea, impairment of mucin exocytosis from goblet cells, e.g. by deletion of vesicle-associated membrane protein 8 (VAMP8), leads to heightened host susceptibility to enteric pathogens *Citrobacter rodentium* and *Entamoeba histolytica* (37, 38). Although the function of goblet cell secretory granules in Muc2^-/-^ mice has not been precisely described, a “flush out” strategy may be ineffective in Muc2^-/-^ mice given the absence of Muc2. Altered mucus homeostasis could lead to goblet cell dysfunction, a phenotype that may increase *L. monocytogenes’* access to its host receptor E-cadherin (39). Indirect consequences of the absence of Muc2, including changes in the composition of the microbiota (16) and elevated basal colonic inflammation (15), may also contribute to the susceptibility of Muc2^-/-^ mice to oral infection with *L. monocytogenes*.

Besides Muc2’s critical physical role in the structure and function of intestinal mucus, mucin glycans can serve as sources of nutrition for pathogens and are important regulators of bacterial pathogenicity (40–42). For example, O-glycans on MUC5AC can suppress the virulence of *Pseudomonas aeruginosa*, facilitating its clearance in a porcine burn wound model (41). The absence of such inter-kingdom regulatory signals in Muc2^-/-^ mice might modulate the *L. monocytogenes* virulence gene program and thus impact *L. monocytogenes* systemic dissemination.

Kumar *et al*. recently reported that Muc2^-/-^ mice have elevated basal levels of systemic inflammation and circulatory iron, conditions that may promote growth of disseminated bacteria and were associated with increased susceptibility to i.p. challenge with LPS (19). This suggested that animals with whole body knockouts of Muc2 have systemic defects in their response to PAMPs. However, such changes did not appear to alter the capacity of Muc2^-/-^ mice to control systemic infection caused by i.p. administration of *L. monocytogenes*, as pathogen burdens were similar after this route of infection in WT and Muc2^-/-^ mice. Further work will be required to decipher the potential extra-intestinal contributions of Muc2 to host defense against pathogens, which may be challenge- and tissue-specific.

The structure of the mucus layer varies in different regions of the intestinal tract; the small intestine is covered by a thin, porous mucus layer while the colon is heavily coated with two layers of Muc2-containing mucus (1). We found that the effects of the absence of Muc2 were particularly marked in the distal colon, where there was a drastic increase in the burden of intracellular *L. monocytogenes;* there was only a moderate elevation in the pathogen burden in the ileum and proximal colon. Consistent with our observations, in a Muc2^-/-^ mouse model of spontaneous colitis, the distal colon was found to have more neutrophilic infiltrates compared to the proximal colon (15). The distal colon is distinguished by a thick inner mucus layer that is not easily penetrated by microbes (35, 43, 44), and a population of goblet cells that supports the rapid renewal of mucus within this inner layer (45). The severe phenotype we observed in the distal colon supports the idea that this region of the intestine is particularly reliant on Muc2-producing goblet cells for defense.

Muc2 deficiency not only affects the pathogen burden in host tissues, but also alters the pathogen’s population dynamics. Our previous (29) and present observations, in two different strains of mice, are consistent with the hypothesis that *L. monocytogenes* disseminates from the intestinal tract to distal organs using multiple independent routes, a pattern that has been referred to as “episodic spread” (29). Notably, the pathogen’s dissemination pattern was altered in the Muc2^-/-^ mice. The genetic relatedness of the *L. monocytogenes* populations recovered from the colon and distal organs (MLN, spleen, and liver) was significantly higher than observed in WT mice, suggesting that the colon is the source for a larger fraction of organisms that disseminate in Muc2^-/-^ versus WT mice. We also observed a marked increase in both the *L. monocytogenes* burden and founding population size in the colon of Muc2^-/-^ mice. Together these observations suggest that Muc2 in the colon is a critical host restriction factor that ordinarily prevents *L. monocytogenes* colonization and invasion of epithelium. The absence of this physical host barrier enables the colon to become the reservoir for dissemination. Heightened basal colonic inflammation in Muc2^-/-^ mice (15) also may contribute to *L. monocytogenes* proliferation and prolong infection. A deeper understanding of the consequences of and mechanisms by which perturbations of intestinal structure and function modify host defense could provide insight to guide new treatment strategies for disorders where intestinal insults promote systemic disease presentations, such as inflammatory bowel disease.

---

**Figure S1.**
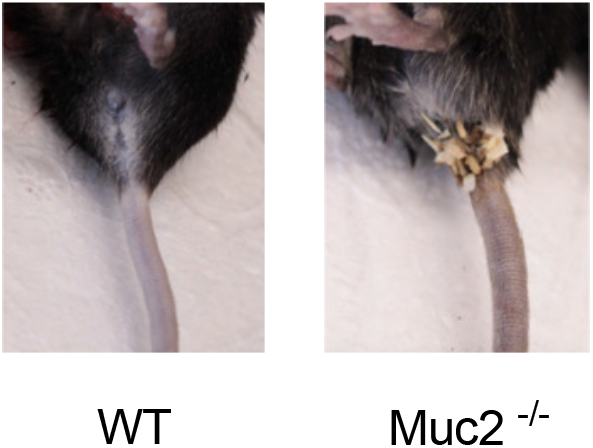
Muc2^-/-^ but not WT mice develop diarrhea after orogastric challenge with *L. monocytogenes*. Muc2^-/-^ mice and WT were challenged via orogastric route with ~3 × 10^9^ CFU of *L. monocytogenes* 10403S InlA^m^. Muc2^-/-^ mice (right) but not WT mice (left) developed diarrhea-like symptoms at 3DPI.

**Figure S2.**
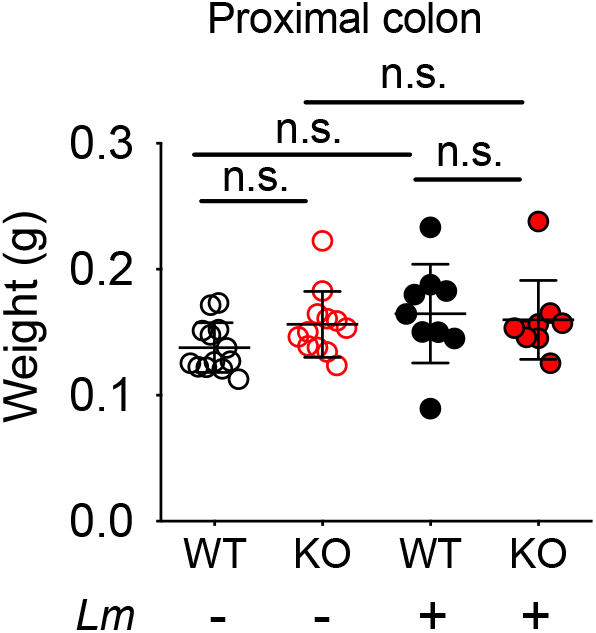
The weight of the proximal colon was not changed by *L. monocytogenes* infection in Muc2^-/-^ mice. Weight of the proximal colon of uninfected and infected Muc2^-/-^ and WT mice at 3 DPI; 9-13 mice/group. ANOVA and Fisher multiple comparison test were used to assess significance (n.s. not significant).

**Figure S3.**
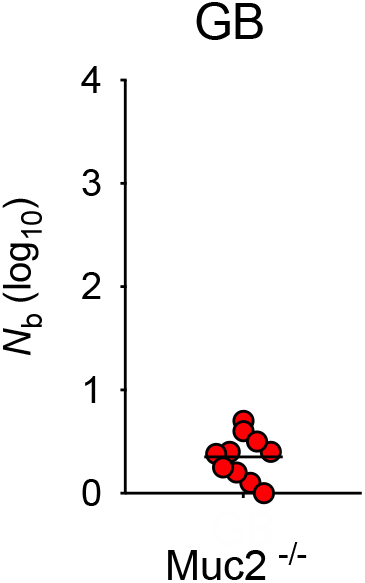
Gallbladder *N*b values in orogastrically infected Muc2^-/-^ mice. *L. monocytogenes* populations that were recovered from bile of Muc2^-/-^ mice at 3 DPI were used for determination of *N*b values; data (n=10) are from three independent experiments.

## Methods

### Bacterial strains and culture conditions

The *Listeria monocytogenes* strain used in this study (Lm-STAMP-200 library, Supplementary table 1) is a barcoded derivative (29) of *L. monocytogenes* 10403S InlA^m^, a strain where internalin A contains two amino acid substitutions that increase its capacity to bind murine E-cadherin (30). For animal studies, aliquots of Lm-STAMP-200 library were cultured in brain heart infusion (BHI; BD Biosciences) broth with chloramphenicol (Cm, 7.5 μg/mL) and streptomycin (Sm, 200 μg/mL) at 37 °C for 3 hours. Bacteria were pelleted by centrifugation (3,000 × g for 10 min), washed twice with 20 ml phosphate-buffered saline (PBS) and diluted to the indicated concentration in PBS for inoculation.

### Animal studies

C57BL/6 Muc2^-/-^ mice used in this study were a gift from Dr. Anna Velcich (13) and were bred at the Harvard Institutes of Medicine animal facility. Mice were genotyped at 3-week-old and Muc2^+/+^ (WT) and Muc2^-/-^ mice were cohoused until 10 to 12-week-old. Gender- and age-matched littermates that were the offspring of heterozygous Muc2^+/-^ breeders were used throughout the study. As observed by other groups (46), a small fraction of Muc2^-/-^ animals (~10 %) in our colony developed spontaneous colon prolapse by 10-12 weeks of age; these mice were euthanized and excluded from the study. For orogastric infection studies, WT and Muc2^-/-^ littermates were fasted for 8 hours prior to infection and orally inoculated with ~ 3 × 10^9^ CFUs of barcoded *L. monocytogenes* (200 μL) suspended in a 300 μL mixture of 200 mM CaCO_3_. For intraperitoneal infection studies, WT and Muc2^-/-^ littermates were administered ~1 × 10^5^ CFU of barcoded *L. monocytogenes* (200 μL) in PBS via i.p. injection. Mouse bodyweights were measured prior to inoculation and daily post-inoculation for 3 days. To assess gross intestinal pathology, animals were euthanized 3 DPI and the length and weight of colons were measured. For survival curve analysis after orogastric inoculation, animals were monitored twice per day for 8 days.

To assess bacterial burden, animals were euthanized 3 DPI and organs [ileum (distal one-third of SI), colon, MLN, spleen, liver, and GB] were collected. Colonic tissues were equally divided into two parts and designated as proximal colon and distal colon. Proximal colons, distal colons, and ileums were cut open longitudinally, incubated in DMEM with 100 μg/mL gentamicin for 2 h to kill the extracellular bacteria, transferred to 50 ml conical centrifuge tubes (Corning), washed with 25 mL of PBS five times on a rotator. The intestinal segments, MLN, spleen, and liver were homogenized in sterile PBS using a beat-beater (BioSpec Products); To collect bile, GBs were transferred to Eppendorf tubes containing 1 mL of PBS and ruptured with a 23-gauge needles (Becton Dickinson). For CFU enumeration, all of the homogenates/samples were serially diluted, plated on BHI-Sm plates, and kept at 37 °C for 48 h. For STAMP analysis, homogenates/samples were directly plated on 245 mm × 245 mm square BHI-Sm plates (Corning).

### Histological analysis

Distal colons from WT and Muc2^-/-^ animals 3 DPI were fixed in 4% paraformaldehyde (PFA) for 2 h, transferred to 70% ethanol and kept at 4°C overnight. Samples were embedded in Tissue-Tek OCT solution (Sakura Finetek) and sliced into 10 μM sections using a Leica CM1860 UV cryostat. Samples were stained with hematoxylin and eosin, mounted with Organo/Limonene mounting medium (Sigma) and scanned (200 × magnification) using a Nikon confocal microscope.

### STAMP protocol

Calculation of N_b_ and genetic relatedness was performed as previously described (29, 32). Briefly, *L. monocytogenes* colonies from the indicated organs were washed off of the BHI-Sm plates with PBS. Cells were pelleted and genomic DNA was extracted (Wizard Genomic DNA Purification Kit; Promega) from ~1 × 10^10^ bacteria. The region that harbors the 30-bp barcodes was amplified from genomic DNA using primer PLM30 and primer PLM6-P29 (Supplementary table 1). The PCR products were purified (MinElute; Qiagen) and quantified (Qubit dsDNA HS Assay Kit; Life Technologies). Purified PCR products were combined in equimolar concentrations and sequenced on an Illumina MiSeq (Miseq Reagent Kit V2, 50-cycle; Illumina) using primer PLM49. Reaper-12–340 was used to discard sequence reads with low quality (≤Q30) and to trim the sequence following the barcode (47). The trimmed sequences were clustered with QIIME (version 1.6.0) using pick_otus.py with a sequence similarity threshold of 0.9 (48). N_b_ was then calculated using a custom R script. Genetic distance was estimated using the Cavalli–Sforza chord distance method (49) as described by Abel et al (32). Genetic relatedness is 1 - genetic distance.

### Ethics statement

Animal experiments in this study were carried out in accordance with the NIH Guide for Use and Care of Laboratory animals and were approved by the Brigham and Women’s Hospital IACUC (2016N000416). Mice were euthanized by isoflurane inhalation followed by cervical dislocation.

## Acknowledgments

We thank Dr. Anna Velcich for providing Muc2^-/-^ mice. We thank members of the Waldor laboratory for helpful discussions. T.Z. was supported by a Sarah Elizabeth O’Brien Trust Postdoctoral Fellowship. This work was supported by NIH Grant RO1-AI-042347 (to M.K.W.), and the Howard Hughes Medical Institute (M.K.W.).

