## Supplemental table 1 for "Muc2 mucin limits *Listeria monocytogenes* dissemination and modulates its population dynamics"

Supplementary table 1. Bacterial strains and primers used in this study.

| Primers | 5' to 3' sequence |
|---|---|
| PLM6 | CAAGCAGAAGACGGCATACGAGATCGTGATGTGACTGGAGTTCAGACGTGTGCTCTTCCGATCTTGTCTCATGAGCGGATACA |
| PLM7 | CAAGCAGAAGACGGCATACGAGATACATCGGTGACTGGAGTTCAGACGTGTGCTCTTCCGATCTTGTCTCATGAGCGGATACA |
| PLM8 | CAAGCAGAAGACGGCATACGAGATGCCTAAGTGACTGGAGTTCAGACGTGTGCTCTTCCGATCTTGTCTCATGAGCGGATACA |
| PLM9 | CAAGCAGAAGACGGCATACGAGATTGGTCAGTGACTGGAGTTCAGACGTGTGCTCTTCCGATCTTGTCTCATGAGCGGATACA |
| PLM10 | CAAGCAGAAGACGGCATACGAGATCACTGTGTGACTGGAGTTCAGACGTGTGCTCTTCCGATCTTGTCTCATGAGCGGATACA |
| PLM11 | CAAGCAGAAGACGGCATACGAGATATTGGCGTGACTGGAGTTCAGACGTGTGCTCTTCCGATCTTGTCTCATGAGCGGATACA |
| PLM12 | CAAGCAGAAGACGGCATACGAGATGATCTGGTGACTGGAGTTCAGACGTGTGCTCTTCCGATCTTGTCTCATGAGCGGATACA |
| PLM13 | CAAGCAGAAGACGGCATACGAGATTCAAGTGTGACTGGAGTTCAGACGTGTGCTCTTCCGATCTTGTCTCATGAGCGGATACA |
| PLM14 | CAAGCAGAAGACGGCATACGAGATCTGATCGTGACTGGAGTTCAGACGTGTGCTCTTCCGATCTTGTCTCATGAGCGGATACA |
| PLM15 | CAAGCAGAAGACGGCATACGAGATAAGCTAGTGACTGGAGTTCAGACGTGTGCTCTTCCGATCTTGTCTCATGAGCGGATACA |
| PLM16 | CAAGCAGAAGACGGCATACGAGATGTAGCCGTGACTGGAGTTCAGACGTGTGCTCTTCCGATCTTGTCTCATGAGCGGATACA |
| PLM17 | CAAGCAGAAGACGGCATACGAGATTACAAGGTGACTGGAGTTCAGACGTGTGCTCTTCCGATCTTGTCTCATGAGCGGATACA |
| PLM18 | CAAGCAGAAGACGGCATACGAGATTGTTGACTGTGACTGGAGTTCAGACGTGTGCTCTTCCGATCTTGTCTCATGAGCGGATACA |
| PLM19 | CAAGCAGAAGACGGCATACGAGATACGGAACTGTGACTGGAGTTCAGACGTGTGCTCTTCCGATCTTGTCTCATGAGCGGATACA |
| PLM20 | CAAGCAGAAGACGGCATACGAGATTCTGACATGTGACTGGAGTTCAGACGTGTGCTCTTCCGATCTTGTCTCATGAGCGGATACA |
| PLM21 | CAAGCAGAAGACGGCATACGAGATCGGGACGGGTGACTGGAGTTCAGACGTGTGCTCTTCCGATCTTGTCTCATGAGCGGATACA |
| PLM22 | CAAGCAGAAGACGGCATACGAGATGTGCGGACGTGACTGGAGTTCAGACGTGTGCTCTTCCGATCTTGTCTCATGAGCGGATACA |
| PLM23 | CAAGCAGAAGACGGCATACGAGATCGTTTCACGTGACTGGAGTTCAGACGTGTGCTCTTCCGATCTTGTCTCATGAGCGGATACA |
| PLM24 | CAAGCAGAAGACGGCATACGAGATAAGGCCACGTGACTGGAGTTCAGACGTGTGCTCTTCCGATCTTGTCTCATGAGCGGATACA |
| PLM25 | CAAGCAGAAGACGGCATACGAGATTCCGAAACGTGACTGGAGTTCAGACGTGTGCTCTTCCGATCTTGTCTCATGAGCGGATACA |
| PLM26 | CAAGCAGAAGACGGCATACGAGATTACGTACGGTGACTGGAGTTCAGACGTGTGCTCTTCCGATCTTGTCTCATGAGCGGATACA |
| PLM27 | CAAGCAGAAGACGGCATACGAGATATCCACTCGTGACTGGAGTTCAGACGTGTGCTCTTCCGATCTTGTCTCATGAGCGGATACA |
| PLM28 | CAAGCAGAAGACGGCATACGAGATATATCAGTGTGACTGGAGTTCAGACGTGTGCTCTTCCGATCTTGTCTCATGAGCGGATACA |
| PLM29 | CAAGCAGAAGACGGCATACGAGATAAAGGAATGTGACTGGAGTTCAGACGTGTGCTCTTCCGATCTTGTCTCATGAGCGGATACA |
| PLM30 | AATGATACGGCGACCACCGAGATCTACACTCTTTCCCTACACGACGCTCTTCCGATCTTGTAAAACGACGGCCAGT |
| PLM49 | ACGCTCTTCCGATCTTGTAAAACGACGGCCAGT |

Bacterial strains

| Strain | Description | Reference |
|---|---|---|
| Lm-STAMP-200 library | barcoded *L. monocytogenes* 10403S InlA$^{m}$ library | (1) |
